# Territory proximity influences aggression and modulates the relationship between social dominance and oxidative stress in a highly social African cichlid fish

**DOI:** 10.64898/2025.12.03.692096

**Authors:** Olivia D. K. Buzinski, Tyler W. Beyett, Zachary D. Hager, Ezekiel T. Maes, Ryan Wong, Peter D. Dijkstra

## Abstract

High social rank has many benefits, including priority access to mates and resources. However, maintaining high rank or social dominance can be physiologically costly, particularly in species that engage in agonistic competition to maintain their dominance status. Previous studies have rarely manipulated the intensity of agonistic competition in controlled settings to isolate the cost of social dominance. We utilized males of the cichlid fish *Astatotilapia burtoni* to investigate how the degree of territoriality influences patterns of oxidative stress. In this species, high-rank males compete for territories to attract mates. We housed males individually with a defendable structure and used clear tank dividers to allow visual access to a size-matched male neighbor housed in an identical setup to manipulate territoriality. Defendable structures were placed either close to the divider (proximal treatment) or further away (distal treatment). We found that males in the distal treatment expressed higher levels of territorial aggression than those in the proximal treatment. While males in the distal treatment did not display higher levels of oxidative stress, territory proximity influenced the relationships between physiological markers of dominance (i.e., testosterone, gonad size) and oxidative balance. Although our work is not consistent with higher competition increasing the physiological cost of high rank, it suggests that the level of competition influences the regulation of oxidative balance relative to social dominance characteristics. We suggest that manipulating the degree of territoriality in a controlled setting provides a unique opportunity to further our understanding of how individuals manage the trade-offs associated with high rank.

**Summary Statement:** Manipulating the amount of space between territories results in variability in the degree of territorial aggression and the interaction of physiological markers of social dominance and oxidative stress.

## Introduction

Many social species form dominance hierarchies, whereby high-rank individuals have primary access to resources and mates relative to low-rank individuals (Snyder-Mackler et al., 2020; Tibbetts et al., 2022). High-rank males often have an upregulated reproductive axis (hypothalamic-pituitary-gonadal (HPG) axis) as indicated by elevated levels of circulating androgens, high levels of aggression, large gonads and elaborate secondary sex characteristics. In contrast, low-rank males exhibit comparatively lower expression of secondary sex characteristics and are often reproductively inactive (Alward et al., 2020; Sapolsky, 2005; Sperry et al., 2010). Defending high rank or social dominance is metabolically expensive, and identifying the nature of these costs is essential to address questions related to life history variation, such as what factors limit dominance tenure or how social dominance influences investment in other traits such as lifespan (Dowling & Simmons, 2009; Monaghan et al., 2009; Ord, 2021; Speakman & Garratt, 2014).

One potential cost of social dominance is oxidative stress, which occurs when the production of reactive oxygen species (ROS) overwhelms the antioxidant capacity. Oxidative stress can result in cellular oxidative damage and negatively affect organismal performance (Birnie-Gauvin et al., 2017; Bize et al., 2008; Metcalfe & Alonso-Alvarez, 2010). Social competition-induced oxidative stress may therefore constrain investment in other life history traits, such as somatic maintenance or mating success (Monaghan et al., 2009; Selman et al., 2012). However, the evidence that social dominance causes higher levels of oxidative stress is mixed. For example, Beaulieu et al. (2014) found that high-rank male mandrills (*Mandrillus sphinx*) engaged in intense social competition during the breeding season (a period of hierarchical instability) exhibited elevated levels of oxidative damage. In another example, high-rank, harem-holding male Seba’s short-tailed bats (*Carollia perspicillata*) and lower-rank sneaker males exhibited similar levels of circulating lipid oxidative damage. However, males with harems had higher levels of circulating antioxidant markers than sneaker males, indicating that while aggressive competition presents a physiological challenge, individuals may mitigate potential oxidative insults by upregulating antioxidant defenses (Fasel et al., 2017; Fernandez et al., 2014). Though many studies have investigated the relationship between social rank and patterns of oxidative stress, few studies have isolated the effect of aggressive competition on physiological stress markers (Pryke et al., 2007), and even fewer have explicitly linked it to patterns of oxidative stress (Dijkstra et al., 2011; Garratt & Brooks, 2015)

The cichlid fish *Astatotilapia burtoni* form social hierarchies in which dominant males maintain territory as a site for reproduction, express bright coloration, have larger gonads, and engage in aggressive competition with neighboring males; subordinate males are typically drab in color and shoal with females (Fernald & Maruska, 2012). Previous studies have shown that oxidative stress levels in dominant males can surpass those of subordinate males (Border et al., 2019, 2021; Fialkowski et al., 2021). The degree of territoriality expressed by dominant males is influenced by the hierarchical stability and composition of the group (Dijkstra et al., 2022; Maruska et al., 2022), and these attributes of the social environment are known modulators of status-based patterns of oxidative stress in this species (Border et al., 2021; Fialkowski et al., 2021). For example, Border et al. (2021) found that hierarchical instability resulted in increased aggression levels in dominant males relative to those in stable conditions; surprisingly, however, there was no significant difference between social states in plasma oxidative damage measurements (d-ROMs). These findings contradict the previously mentioned study in mandrills where social instability and increased aggression was correlated with oxidative damage, suggesting a complex relationship between aggression and patterns of oxidative stress.

Studies using *A. burtoni* that were conducted using large groups of both males and females present complex social dynamics and sexual interactions that may confound the link between the degree of territoriality, social status and oxidative stress patterns (Border et al., 2019, 2021; Fialkowski et al., 2021). To isolate the effect of territoriality on patterns of oxidative stress, we housed individual *A. burtoni* males individually in tank compartments, each provided with one defendable structure and visual and chemical access to an adjacent compartment. We manipulated the proximity of the structures to the adjacent compartment to promote variation in the degree of territoriality expressed by experimental males. This allows us to gain further insight into how patterns of oxidative stress may be influenced by the degree of territoriality. We predicted that 1) increased territory proximity would heighten aggression and 2) more intense territorial aggression would increase oxidative stress as indicated by more oxidative damage and reduced antioxidant availability.

## Methods

### Animals and Housing

All fish were male *Astatotilapia burtoni* (approx. 8-12 months) bred from a laboratory population originating from Lake Tanganyika (Fernald & Hirata, 1977). Fish were taken from 400 L mixed-sex tanks maintained at 28 °C and kept on a 12 hr: 12 hr light:dark cycle with 10-minute dusk and dawn periods. During experimentation, all fish were fed cichlid flakes and pellets (Omega Sea LLC, Earth City, MO, USA) each morning. Tanks were supplied with gravel substrate and continuous waterflow; water filtration occurred mechanically and biologically. All procedures were approved by Central Michigan University’s Institutional Animal Care and Use Committee (2021-460).

### Experimental Design

Males lacking visible characteristics of social dominance were selected and measured for standard length (SL) and weight at the beginning of the experiment (average weight = 2.393043 g; average length = 45.63043 mm). Following a previously described protocol (Fialkowski et al., 2021), 110 L tanks were divided into two compartments using clear, perforated screens to allow for visual and chemical communication between males. We ensured that the males in each tank were of a similar size and placed a halved terracotta pot in each compartment to serve as defendable structures referred to as caves. We manipulated the distance between territories by placing caves close to the tank divider (proximal treatment, ∼1.5 cm from divider, ∼ 3 cm between pots) or far from the tank divider (distal treatment, ∼30 cm from divider; ∼ 60 cm between pots; Figure 1A). Caves were placed approximately 2cm away from the back of the tank, allowing males to fully enter their cave. A total of 50 fish were used in this study with a final sample size of 22 fish in the distal treatment and 24 fish in the proximal treatment. Four fish died during the first week of the experiment due to aggressive competition after entering the compartment of their neighboring male though small spaces between the edge of the divider and the side of the tank (three from distal treatment, one from proximal treatment). All tank dividers were secured or confirmed to be secure prior to replacing deceased fish with similarly sized males.

**Figure 1:**
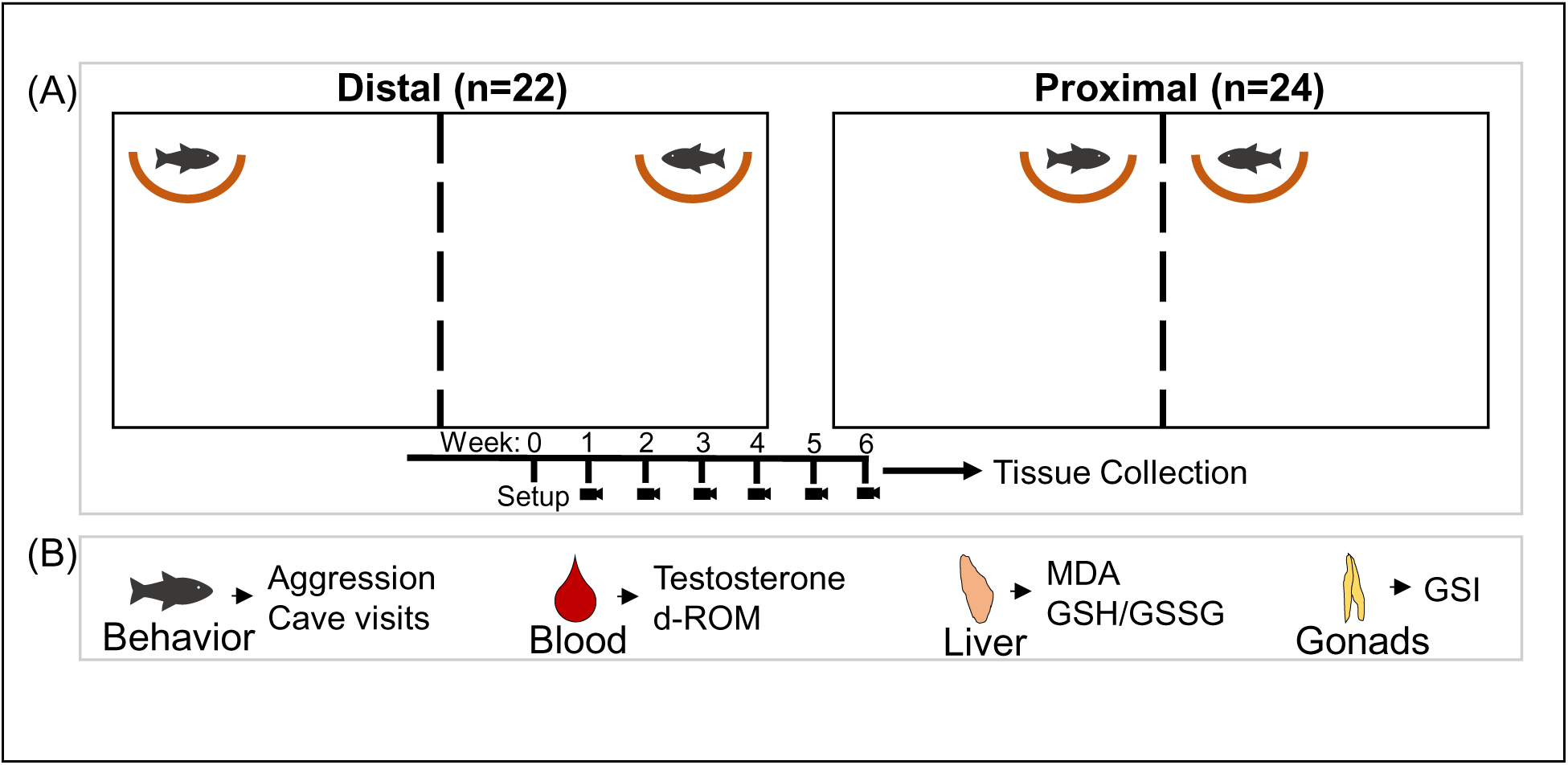
Experimental design. (A) Adult (8-12 months) male *A. burtoni* were paired with a size matched conspecific and housed in neighboring compartments divided using a clear, perorated screen. Caves were placed either far from the divider (distal, n = 22) or close to the divider (proximal, n = 24). Tanks were filmed for 5.5 minutes once per week, for six weeks; after week 6 filming, all fish were sacrificed for tissue collection. (B) Behavior was quantified, and blood, liver, and gonads were collected from each fish for assessing physiological markers of dominance (testosterone, GSI) and oxidative stress (d-ROM, MDA, GSH/GSSG).

### Behavioral Observations

Following a one-week settling period, tanks were filmed in the morning with a Canon EOS 70D camera (Canon U.S.A., Inc., Melville, NY, USA) for 5.5 minutes once per week for six weeks (Figure 1B). One tank (two fish) was included in each video and behavior was quantified for each fish by a single observer for a 5-minute period within each recording, excluding the initial and final 15 seconds of each video while observers were in front of tanks to work with the camera. Cave visits (male enters their pot) were quantified and aggressive behaviors (lateral displays, border displays, and chases) directed toward the neighboring male were quantified as described previously (Fialkowski et al., 2021). Total aggression was calculated by finding the sum of all aggressive behaviors (aggressive displays and chasing) per 5-minute observation period.

### Tissue Preparation

We collected tissue from all males immediately following the filming in week six (Figure 1B). Males were netted and measured for standard length and weight before drawing blood from the caudal vein using heparinized 26-guage butterfly needles (Terumo Medical Corporation, Somerset, NJ, USA); blood samples were kept in heparinized microcentrifuge tubes on ice until they were centrifuged at 4000 x g for 10 minutes to separate the plasma, which was then aliquoted for assays and stored at −20 °C. Next, we sacrificed males via cervical transection prior to collecting gonads which were weighed for the calculation of gonadosomatic index (GSI); livers were collected and weighed, frozen on dry ice, and stored at −20 °C until further processing. For analysis, tissue was homogenized using a D1000 Homogenizer with a 5 mm generator probe (Benchmark Scientific, Sayreville, New Jersey, USA) in a general homogenization buffer (10 µL buffer/mg tissue) consisting of potassium phosphate buffer (50 mM PBS (pH 7.4) and 0.5 mM EDTA) with a Protease Inhibitor Cocktail (Sigma-Aldrich, Inc., P2714, St. Louis, MO, USA) added at a 1:10 dilution.

### Measurements of Oxidative Stress

Homogenized tissue samples were used for various assays to assess markers of oxidative stress (Figure 1B). Liver glutathione was quantified as representative for antioxidant function, malondialdehyde (MDA), a byproduct of lipid peroxidation by ROS, was used as an indicator of liver oxidative stress levels, and reactive oxygen metabolite derivatives (d-ROMs) were quantified in plasma samples as a marker of overall oxidative stress. All assays were run using clear, flat-bottom 96-well plates with the exception of the glutathione assay where black half-area 96-well plates were used; researchers were blinded with respect to treatment.

### Liver Glutathione

Glutathione was quantified in liver homogenate samples (proximal n = 23; distal n = 20) using a DetectX® Glutathione Fluorescent Detection Kit (Arbor Assays, Ann Arbor, MI, USA) per the manufacturer’s protocol previously described by Dijkstra et al. (2024). A Tecan M200 Infinite plate reader (Tecan Group Ltd., Männedorf, Switzerland) was used to find the concentration of free reduced glutathione (fGSH). To find the concentration of oxidized glutathione dimers (GSSG), 25 µL of kit reaction mixture was added to each well followed by a second 15-minute incubation to convert all GSSG in the sample to fGSH before a second fluorescence reading to find the total concentration of reduced glutathione (tGSH) in each sample; GSSG was calculated ((tGSH - fGSH) / 2) to find the ratio of fGSH/GSSG. The average interplate control CV for free glutathione measurements was 10% (two batches), and the average intraplate control CV was 3%. For total glutathione measurements, the average interplate control CV was 8% (two batches), and the average intraplate control CV was 2%.

### Liver Lipid Peroxidation

Malondialdehyde (MDA) concentrations were quantified in 150 µL liver homogenate samples (proximal n = 23; distal n = 22) according to the colorimetric thiobarbituric acid reactive substances (TBARS) assay kit protocol (Cayman Chemical, Ann Arbor, MI, USA). Plates were read using an Epoch2T plate reader (Biotech Instruments, Winooski, VT, USA) to obtain MDA (µM) values. The average intraplate control CV was 13%, an interplate control was not used.

### Circulating d-ROM

To assess overall oxidative stress, 4 µL plasma samples (proximal n = 14; distal n = 19) were processed to measure reactive oxygen metabolite derivatives (d-ROM) following a previously described kit protocol (Diacron, Grosseto, Italy; Border et al., 2019). Absorbance values were measured using the Tecan M200 Infinite plate reader, and concentrations were reported as H_2_0_2_ mg dL^−1^. All samples were run on one well-plate, the average intraplate CV was 14%.

### Circulating Testosterone

Testosterone was selected as a measure of HPG axis activity and quantified in 7 µL plasma samples (proximal n = 14; distal n = 19) using a previously described competitive-binding ELISA kit and protocol (Enzo Biochem, Inc., Farmingdale, NY, USA; Border et al., 2019). Optical density was read using the Tecan M200 Infinite plate reader. Testosterone values were reported as testosterone ng mL^−1^. The interplate control CV was 10% (two batches), the average intraplate control CV was 2%.

### Statistical Analysis

All analyses were completed in R v4.30 (R Core Team, 2020) using the R packages lme4, lmerTest, MASS (Bates et al., 2015), glmmTMB (Brooks et al., 2017), and emmeans for posthoc comparisons (Lenth, 2025). Assay sample sizes varied depending on amount or quality of available sample. GSI ((gonad mass/body mass) * 100) and body condition (weight/(length^3^) * 100) were calculated (Bolger & Connolly, 1989).

To assess the effect of our treatment on the selected HPG axis and oxidative stress markers, we used linear mixed models (LMM) with fish pair as a random effect. For the behavioral analyses, we used generalized mixed models assuming a negative binomial distribution (GLMM). We used fish pair as a random effect when analyzing behavior as a predictor for physiological markers and individual fish ID as a random effect in assessing behavior over time. To evaluate the validity of our models, we examined plots of predicted versus residuals. We used Tukey’s multiple comparison post hoc tests for pairwise comparisons between groups. A significance level (*a*) of 0.05 was used for all tests. We report mean ± SE.

## Results

### Cave Position and Behavior

Total aggression varied significantly over time and across treatments (Table 1). Total aggression significantly decreased over time in the proximal treatment (GLMM, −0.110 ± 0.046, z = −2.406 *p* = 0.016) and increased in the distal treatment (GLMM, 0.169 ± 0.046, z = 3.628, *p* = 0.0003). Post-hoc comparisons showed that total aggression was significantly higher in proximal treatment males relative to males in the distal treatment during the first week, after which point males in the distal treatment expressed higher aggression levels than the proximal treatment males, reaching significance in weeks five and six (Figure 2A). Overall, males in the distal treatment performed significantly more aggressive behaviors during the experiment (GLMM, −0.266 ± 0.118, z = 2.26, *p* = 0.024). Males in the proximal treatment performed more cave visits than males in the distal treatment throughout the experiment (GLMM, 0.639 ± 0.146, z = 4.377, *p* < 0.0001, Figure 2B) but there was no significant interaction effect of treatment and week on the number of cave visits performed (Table 1). There were no significant differences in testosterone and GSI across treatment groups (testosterone: LMM, −2.013 ± 3.889, t = 0.517, *p* = 0.608; GSI: LMM, −0.063 ± 0.038, t = −1.637, *p* = 0.109; Figure 3A)

**Figure 2:**
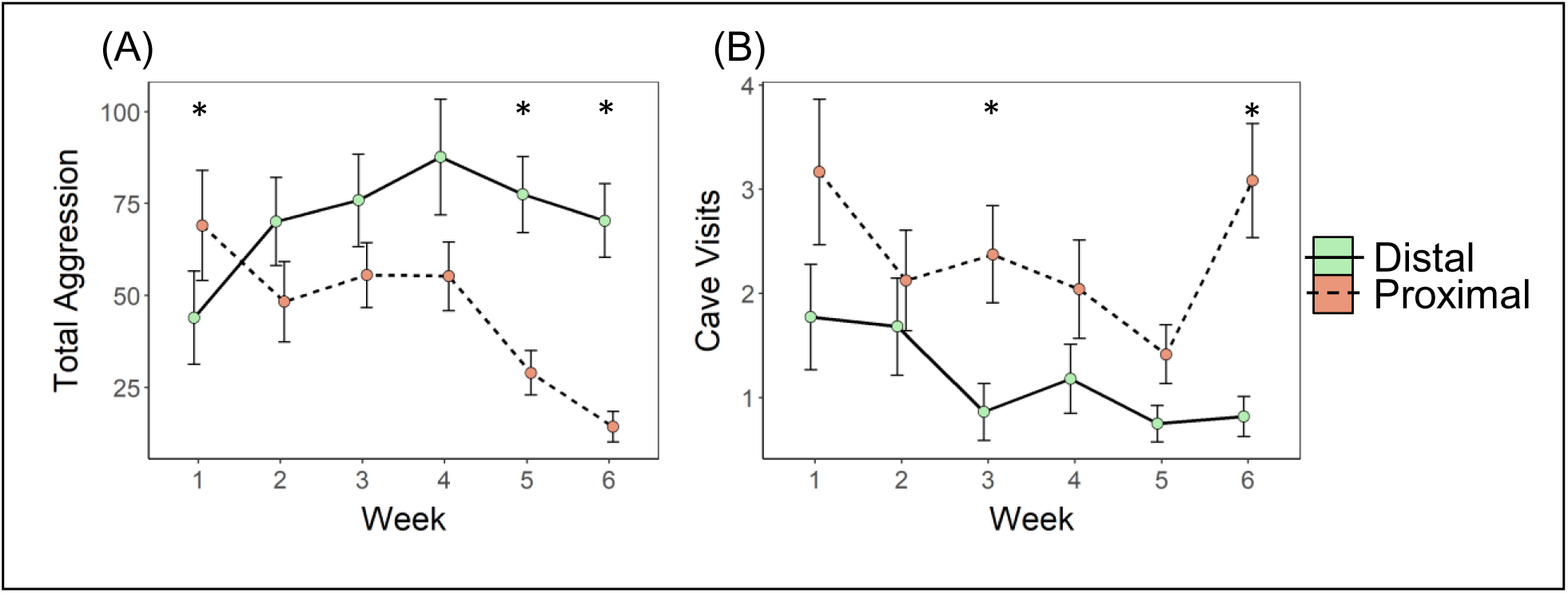
Behavior over time. Group means were calculated for distal treatment (solid line) and proximal treatment (dashed line), error bars represent standard error. (A) Total aggression (sum of aggressive behaviors) was calculated for each observation period and was compared across treatments for each week. (B) The total number of cave visits performed in each observation period was found and compared across treatments for each week (\**p* < 0.05).

**Figure 3:**
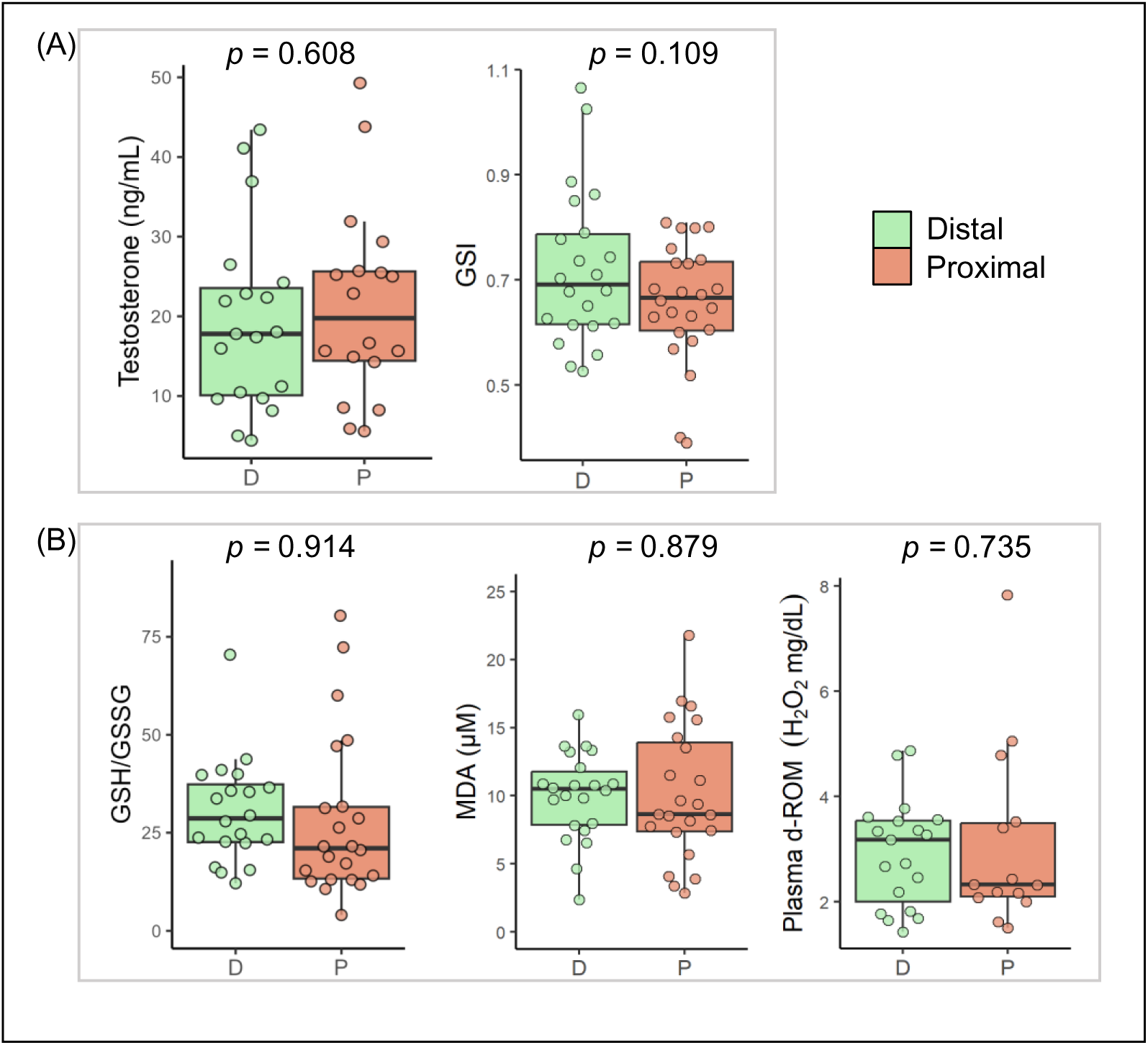
HPG axis and oxidative stress measurements for distal treatment (green) and proximal treatment (orange). (A) Testosterone and gonadosomatic index (GSI) were used as markers of the HPG axis. (B) GSH/GSSG (higher ratio indicates lower oxidative stress levels), MDA (oxidative damage marker), and d-ROM (oxidative damage marker). Error bars enclose data range, excluding outliers.

**Table 1.** Behavior over time. Cave visits and total aggression week (sum of all aggressive behavior counts) was calculated for each fish each week. Generalized mixed models assuming a negative binomial distribution were used to assess treatment and week interaction effects on the behavioral markers, fish ID and tank were included as random effects. *p* < 0.05*, *p* < 0.001*\*\*, p* < 0.0001*\*\*\**

| Marker | Effect | Estimate | Statistic | *p* value |
| --- | --- | --- | --- | --- |
| Total aggression | Week | 0.14718 ± 0.03976 | *z* = 3.701 | **0.000214\*\*** |
|  | Treatment | 0.66412 ± 0.30407 | *z* = 2.184 | **0.028958\*** |
|  | Week x Treatment | -0.29559 ± 0.0583 | *z* = -5.07 | **<0.0001\*\*\*** |
| Cave visits | Week | -0.09531 ± 0.06691 | *z* = -1.424 | 0.154 |
|  | Treatment | 0.42398 ± 0.35959 | *z* = 1.179 | 0.238 |
|  | Week x Treatment | 0.06307 ± 0.0813 | *z* = 0.776 | 0.438 |

### Hypothalamic-Pituitary-Gonadal (HPG) Axis and Oxidative Stress

GSH/GSSG, d-ROM, and MDA measurements did not significantly differ by treatment (GSH/GSSG: LMM, 0.08386 ± 0.06939, t = 1.209, *p* = 0.234; MDA: LMM, 0.228 ± 1.489, t = 0.153, *p* = 0.879; d-ROM: LMM, 0.178 ± 0.510, t = 0.350, *p* = 0.735; Figure 3B). We found a significant interaction effect of treatment and GSI on MDA levels but no significant interaction effect of treatment and GSI on GSH/GSGG or d-ROM (Table 2; Figure 4A). However, we found that testosterone and treatment did have a significant interaction effect on GSH/GSSG but no effect on MDA and d-ROM (Table 2; Figure 4B).

**Figure 4:**
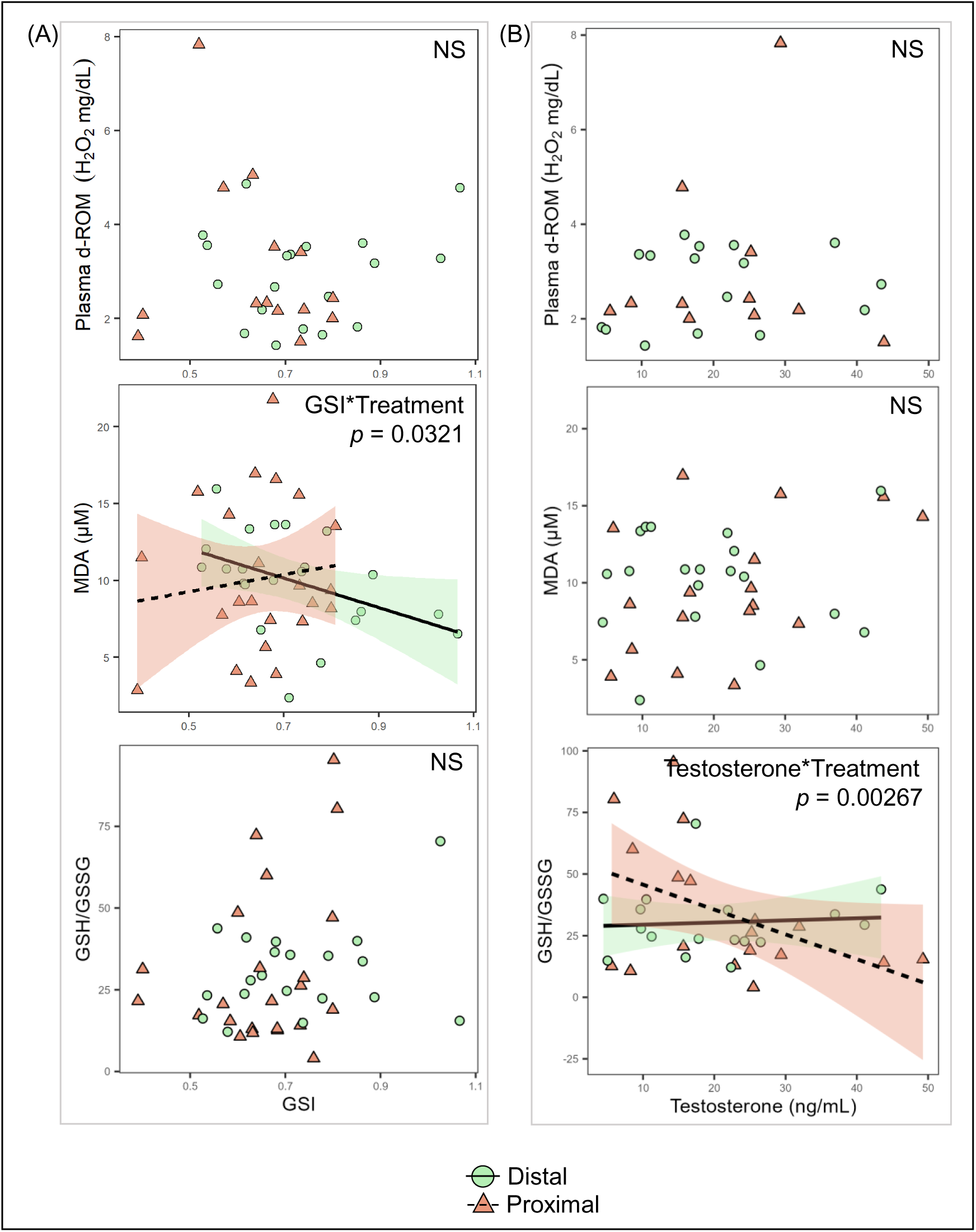
HPG axis and oxidative stress marker interactions. Treatment interaction effects are presented with separate lines for the distal treatment (solid line, circle datapoints) and proximal treatment (dashed line, triangle datapoints). (A) oxidative stress markers shown in response to GSI scores. (B) oxidative stress markers shown in response to testosterone measurements. Shaded regions represent 95% confidence intervals.

**Table 2.** HPG axis effects on oxidative stress. Linear mixed models were used to assess interaction effects of HPG axis markers (testosterone and GSI) and treatment on the selected markers of oxidative stress (GSH/GSSG, MDA, d-ROM). Tank number was included as a random effect.. *p* < 0.05*, *p* < 0.001*\*\*, p* < 0.0001*\*\*\**

| Marker | Effect | Estimate | df | t value | p value |
| --- | --- | --- | --- | --- | --- |
| GSH/GSSG | GSI | 16.45 ± 26.97 | 32.49 | 0.610 | 0.546 |
|  | Treatment | -38.53 ± 29.53 | 29.90 | -1.305 | 0.202 |
|  | GSI x Treatment | 61.63 ± 42.04 | 26.42 | 1.466 | 0.154 |
| GSH/GSSG | Testosterone | 0.0324 ± 0.3317 | 22.5708 | 0.098 | 0.92298 |
|  | Treatment | 35.8406 ± 12.1363 | 34.5669 | 2.953 | 0.00562 |
|  | Testosterone x Treatment | -1.4988 ± 0.4402 | 20.9720 | -3.405 | 0.00267 |
| MDA (µM) | GSI | -13.647 ± 3.88 | 25.147 | -3.518 | 0.00168 |
|  | Treatment | -10.049 ± 4.555 | 30.848 | -2.206 | 0.03496 |
|  | GSI x Treatment | 14.398 ± 6.344 | 25.123 | 2.27 | 0.03208 |
| MDA (µM) | Testosterone | -0.0065 ± 0.0722 | 26.9456 | -0.090 | 0.929 |
|  | Treatment | -3.7783 ± 2.5714 | 31.0494 | -1.469 | 0.152 |
|  | Testosterone x Treatment | 0.1475 ± 0.1011 | 28.3698 | 1.459 | 0.156 |
| d-ROM (H<sub>2</sub>O<sub>2</sub> mg/dL) | GSI | 1.352 ± 2.062 | 19.462 | 0.656 | 0.52 |
|  | Treatment | 3.19 ± 2.425 | 21.773 | 1.316 | 0.202 |
|  | GSI x Treatment | -4.451 ± 3.482 | 19.221 | -1.278 | 0.216 |
| d-ROM (H<sub>2</sub>O<sub>2</sub> mg/dL) | Testosterone | 0.03684 ± 0.0254 | 10.9601 | 1.453 | 0.1744 |
|  | Treatment | 1.2803 ± 1.2051 | 22.7770 | 1.062 | 0.2992 |
|  | Testosterone x Treatment | -0.0438 ± 0.0445 | 16.5092 | -0.985 | 0.3389 |

## Discussion

Oxidative stress has been indicated as a potential cost of social dominance, though pinpointing how specific aspects of social dominance drive variation in oxidative stress has proven to be difficult given the contribution of numerous factors (e.g., environment, season, individual differences) to overall oxidative status (Cram et al., 2015; Fialkowski et al., 2021; Georgiev et al., 2015). Here, we manipulated the level of social competition by placing defendable structures symmetrically on either side of a clear divider either close together (proximal treatment) or far apart (distal treatment) and housed size-matched males in isolation on both sides of the divider. We found that after the first week of the experiment, males in the distal treatment expressed higher levels of aggression relative to males in the proximal treatment, reaching significance in weeks five and six. During weeks five and six, average total aggression levels in the distal treatment were about three times higher than what was observed in proximal treatment males. We predicted that males with experimentally heightened aggression would exhibit higher levels of oxidative stress, but we failed to detect an effect of cave distance treatment on oxidative stress. Interestingly, we found that the relationships between physiological markers of social dominance (i.e., testosterone, gonad size) and oxidative balance were influenced by treatment.

The increased level of aggression in the distal treatment relative to the proximal treatment contrasted with our prediction. However, our findings are consistent with the notion that ‘dear enemy’ effects (i.e., the phenomenon that territory owners exhibit diminished responses to familiar territorial neighbors versus strangers) might be stronger when neighbors are in close proximity to each other, a result of more efficient information gathering from both agonistic and non-agonistic interactions or events (Carazo et al., 2008; Temeles, 1994). Another explanation could be that males in the distal treatment perceived their territories as larger with more ‘resource value’ and were consequently more motivated to defend it (Breau & Grant, 2002; Leese & Blatt, 2021). For example, increased territory size is correlated with heightened aggression in cichlids and other animals species (Mumby & Wabnitz, 2002; Sturmbauer et al., 2008; Watson & Miller, 1971). Regardless of the underlying cognitive and neuroendocrine mechanisms, our experiment induced substantial differences in the intensity of territorial defense by simply varying cave distance between individually housed males. Contrary to our prediction, the higher rate of aggressiveness in the distal treatment did not appear to induce higher levels of oxidative stress compared to those that were housed in the proximal treatment. Our findings here suggest that males can manage oxidative stress under varying levels of social competition, as has been suggested in *A. burtoni* and other species (Garratt et al., 2012). For example, in a previous report from our group, we found that *A. burtoni* males that were allowed to ascend from subordinate to dominant status upregulated enzymatic antioxidants within a few days after social ascent, presumably to cope with the metabolic challenges of competition and upregulation of the reproductive system (Fialkowski et al., 2021). The present study adds to a growing body of literature indicating that metabolically challenging behaviors do not necessarily lead to increased expression of physiological stress markers due to protective or plastic mechanisms or behaviors (Gilmour et al., 2025; Sawecki & Dijkstra, 2022; Van Dievel et al., 2016).

We found that the relationship between testosterone levels and aggression at the end of the experiment (week six, prior to tissue collection) differed by treatment. In dominant male *A. burtoni*, testosterone levels are typically positively associated with aggression, however this trend was only present in males of the distal treatment in our study. In other words, while there was no treatment effect on testosterone, the typical pattern of aggression levels increasing with testosterone was only observed when caves were placed distally while males in the proximal treatment did not exhibit higher aggression levels as testosterone increased. Since a positive link between testosterone levels and relative gonad size could be an indicator of increased regulation of the reproductive system and competitive behavior (Maruska & Fernald, 2010), our findings provide further support for males exhibiting increased territoriality in the distal treatment. We note that there was considerable overlap in both testosterone levels and relative gonad size (GSI) across treatments, which presented an opportunity to investigate how individuals that exhibited similar testosterone levels and reproductive investment but dramatically different levels of competitive effort differ in oxidative balance (Alonso-Alvarez et al., 2007; Dijkstra et al., 2012).

We found that cave treatment influenced how circulating testosterone levels were correlated with GSH/GSSG as well as with plasma oxidative damage (d-ROM). These findings are consistent with the notion that androgens influence ROS production and antioxidant responses (Holmes et al., 2013; Lu et al., 2010). The fact that cave distance treatment (which yielded different levels of aggression) modulated the link between testosterone and oxidative stress could suggest that such androgenic regulation of oxidative balance may be dependent on the metabolic state of the animal, as has been found in previous studies (Holmes et al., 2013; Snyder et al., 2018). Further, we found a treatment effect on the relationship between GSI and liver lipid peroxidation levels (MDA); we detected a significant negative relationship between GSI and liver MDA in the distal treatment, while a non-significant positive relationship was found in the proximal treatment. However, inferring causality for such patterns requires direct manipulation of androgenic signaling or gonadal growth using pharmacology (Huffman et al., 2012) or mutant males with disrupted androgen receptors (Alward et al., 2020). Nevertheless, the observed treatment-dependent relationships between physiological dominance characteristics (i.e., testosterone, gonad size) and oxidative stress suggest that cave distance (or level of competition) influences the regulation of oxidative balance under varying levels of reproductive investment, perhaps by impacting potential trade-offs between reproductive activity and management of oxidative stress. In other words, our study provides experimental support that the level of competition impacts the link of oxidative stress and the degree of investment in reproduction and territoriality, supporting the idea that oxidative stress may play a role in mediating trade-offs between competing life history traits (Dowling & Simmons, 2009; Monaghan et al., 2009; Speakman & Garratt, 2014).

Higher levels of competitive behaviors did not result in increased oxidative stress, but territory proximity did influence the relationship between physiological dominance characteristics (i.e., testosterone, gonad size) and several markers of oxidative stress. Our results suggest that the social environment influences how high-ranking individuals cope with the physiological costs associated with dominance. We suggest that manipulating the degree of territoriality in a controlled setting with only visual access to one conspecific provides an opportunity to gain insight into the cost of aggression and physiological dominance phenotypes in the absence of potentially confounding factors (e.g., injury, sexual opportunity, hierarchical instability). Our study adds to a growing body of literature showing that organisms have sophisticated mechanisms to cope with potential oxidative insults during challenging life history events (Birnie-Gauvin et al., 2017; Costantini, 2008; Giraud-Billoud et al., 2024). Insights into these adaptations may reveal specific physiological trade-offs or costs that do not simply result from higher levels of ROS or oxidative stress. For example, mounting an antioxidant response may have unwanted pleiotropic effects or cellular adaptation to oxidative stress may be metabolically expense. Future work should therefore clarify specific mechanisms underlying the regulation of oxidative balance under varying levels of competition.

## Acknowledgements

We thank members of the Dijkstra lab for providing helpful comments on earlier drafts of the manuscript, and Cooper Borders for his contribution to data collection.

## Data and resource availability

Data will be made available upon acceptance of this article.

